# Cellular production of a *de novo* membrane cytochrome

**DOI:** 10.1101/2022.12.06.519282

**Authors:** Benjamin J Hardy, Alvaro Martin Hermosilla, Dinesh K Chinthapalli, Carol V Robinson, JL Ross Anderson, Paul Curnow

## Abstract

Heme-containing integral membrane proteins are at the heart of many bioenergetic complexes and electron transport chains. The importance of these electron relay hubs across biology has inspired the design of *de novo* proteins that recreate their core features within robust, versatile and tractable protein folds. To this end, we report here the computational design and in-cell production of a minimal diheme membrane cytochrome which successfully integrates into the cellular membrane of live bacteria. This synthetic construct emulates a four-helix bundle found in modern respiratory complexes but has no sequence homology to any polypeptide sequence found in nature. The two *b*-type hemes, which appear to be recruited from the endogenous heme pool, have distinct split redox potentials with values close to those of natural membrane-spanning cytochromes. The purified protein can engage in rapid biomimetic electron transport with small molecules, with other redox proteins, and with biologically-relevant diffusive electron carriers. We thus report an artificial membrane metalloprotein with the potential to serve as a functional module in electron transfer pathways in both synthetic protocells and living systems.

## INTRODUCTION

Integral membrane proteins underpin bioenergetic reactions across all domains of life. This functionally-diverse group includes the ubiquitous membrane-embedded hemoproteins that are active redox centres in respiratory and photosynthetic electron transport chains. The heme cofactors coordinated by these proteins provide conduits for electron transfer that can link redox events on opposite sides of the membrane, allow the transfer of electrons to membrane-embedded active sites, or connect membrane quinol/quinone (Q) pools to extramembrane partners for redox catalysis and proton pumping. Classical examples are the mitochondrial and bacterial succinate dehydrogenase (Complex II), mitochondrial cytochrome *bc*_*1*_ (Complex III), heme-copper oxidases such as mitochondrial Complex IV or bacterial *bo*_*3*_ complex, and numerous other proteins of bacterial respiration including fumarate reductase, nitrate reductase and formate dehydrogenase^1-7^. Parallel to the ongoing study of these bioenergetic proteins, there remains an interest in developing non-natural transmembrane cytochromes that can reproduce the essential features of their natural counterparts. The design and engineering of such minimal bioenergetic proteins reveals the critical assembly pathways and general operating principles of heme-containing electron transport complexes, and enables the implementation of their redox and catalytic functions in synthetic systems^8,9^. The fullest realisation of this concept would be to directly integrate these artificial proteins into the respiratory and metabolic pathways of living cells, rerouting electrons to or from the Q-pool to drive biochemical reactions.

Many respiratory cytochromes contain two molecules of *b*-type heme in their transmembrane domains. A long-standing goal in cytochrome design is therefore the creation of a minimal protein architecture that can house two *b*-type hemes at an appropriate distance and orientation for electron transport within, and across, a cellular lipid membrane. Previous work has largely focused on designing individual peptides that can self-assemble into homocomplexes to bind natural and non-natural porphyrins *in vitro*^10-14^. However, these chemically-synthesised peptides cannot be easily incorporated into cellular membranes and are therefore not amenable to biological applications. When designs have been genetically-encoded for biosynthesis via recombinant expression they are either not recovered directly from cell membranes^12,15^ or unable to complex heme *in vivo*^16^, and so rely instead on *in vitro* reconstitution and subsequent heme loading. Until now, artificial membrane cytochromes that are fully compatible with living cells have remained elusive.

A recent study^17^ described the recombinant production of a *de novo* soluble 4-helix bundle, known as 4D2, that coordinates two bis-His ligated *b*-type hemes when produced in *Escherichia (E*.*) coli*. This designed construct (PDB ID 7AH0) was based upon the mutual structure of the heme-binding centres from the respiratory *bc1* and photosynthetic *b*_*6*_*f* complexes, both of which feature four transmembrane alpha helices in direct contact with the heme cofactor^2,18^. The conserved structure of these four helices can be approximated as an antiparallel four-stranded coiled-coil, and this fold is generally accessible to *de novo* metalloprotein design^18,19^. The designed 4D2 protein based upon this fold supports the binding of two *b*-type hemes in closer proximity than is observed in the natural cytochromes *b* (6 Å in 4D2 *vs* 12-13 Å in *bc*_1_), as a consequence of placing the heme-ligating histidines in a symmetrical fashion on alternating helices. The advantage of such a short interheme distance is the potential for exceptionally rapid electron transfer between the cofactors.

Here, we report the successful conversion of water-soluble 4D2 into a functional integral membrane hemoprotein that is fully compatible with living cells. Computational redesign of the 4D2 surface introduces hydrophobic residues that are compatible with protein insertion into a lipid bilayer. The resulting construct, which we call Cytochrome *b*X (CytbX), can be produced by recombinant *E. coli*, is targeted to the cell membrane, and can be purified from membrane fractions in complex with two endogenous *b*-type hemes. This *de novo* transmembrane cytochrome can take part in chemical and biologically-relevant electron transfer reactions, including reconstituted electron transport pathways. These results establish a basis for the future design of customised membrane-embedded metalloproteins with broad application.

## MATERIALS AND METHODS

### Computational design

Residues of the 4D2 bundle forming interhelical knobs-into-holes interactions were identified with SOCKET^20^. All design was performed using Rosetta versions 3.12 with the *franklin2019* energy function^21,22^, and full design protocols are included as Supplementary Data. Briefly, surface residues for mutation were explicitly specified from PDB deposition 7AH0 and restricted to amino acids with hydrophobic and small polar sidechains (FAILVWGST). An amino acid composition score term was included to limit the total count of aromatic groups. A total of 18,400 decoys were generated using the FastDesign and FastRelax movers. The transmembrane topology of selected designs was assessed with TMHMM2.0^23^ or DeepTMHMM^24^and membrane embedding was modeled with PPM 3.0^25^. The sequences of characterised designs are provided as Supplementary Figures S1 and S2, and details of subsequent molecular dynamics simulations are provided as Supplementary Information.

### Molecular biology

CytbX was ordered as a synthetic gene (Twist Biosciences) that was codon-optimised for expression in *E. coli* and included a C-terminal V5 antibody epitope and His_10_ tag. This was cloned into the NcoI/XhoI sites of pET28 and was subsequently used as the basis of a GFP fusion protein in the same vector. The gene products of the His_10_ tag (CytbX) and GFP fusion (CytbX-GFP) have molecular weights of 15,607 Da and 43,333 Da, respectively. Plasmids were transformed into the BL21 derivative C43(DE3) for recombinant expression^26^.

### Protein expression

CytbX was expressed and purified essentially as described previously^16,27^. Expression cultures were 1L LB medium in 2.5 L baffled flasks (Tunair). Cultures were inoculated from an overnight starter at 1:100 dilution and grown at 37°C, 250 rpm to an optical density at 600 nm (OD600) of 0.9. Expression was induced with 0.1 mM IPTG and at the point of induction cultures were supplemented with 25 mg/L of the heme precursor 8-aminolevulinic acid (ALA). Induction was for 2h at 37°C with continued agitation at 250 rpm. Cells were harvested at 5000 x g for 30 mins and the resulting cell pellet stored at -20°C.

### Isolation of cell membranes and protein solubilisation

The thawed cell pellet was resuspended in 100 ml 1x phosphate-buffered saline (PBS) and passed through a continuous flow cell disruptor (Constant Systems) at 25 KPSI. Unbroken cells and inclusions were removed by centrifugation at 10,000 x g, 10 min. Cell membranes were obtained from this clarified lysate by ultracentrifugation at 180,000 x g for 1h. The membrane pellet was resuspended in 40 ml Membrane Buffer (50 mM sodium phosphate buffer at pH 7.4, 150 mM NaCl, 5% v/v Glycerol). After 10 passes in a handheld glass homogeniser the surfactant 5-cyclohexyl-1-pentyl--D-maltopyranoside (Cymal-5) was introduced at 2.4 % w/v. Membranes were solubilised for 1h at room temperature with gentle agitation on a rocker-roller. Solubilized membranes were recovered after ultracentrifugation at 180,000 x g for 1h. Imidazole was added at 20 mM prior to immobilised metal affinity chromatography.

### Protein purification

Solubilized His-tagged CytbX was purified on a 1 ml Ni-NTA column (Cytiva HisTrap). The Purification Puffer (PB) was Membrane Buffer plus 0.24% w/v Cymal-5. The column was equilibrated in at least 5 column volumes of PB with 20 mM imidazole. After protein binding at a flow rate of 1 ml/min the column was washed with 30 column volumes of PB containing 75mM imidazole. Purified CytbX was eluted with 2.5 column volumes of 0.5 M imidazole in PB and the imidazole was immediately removed with a P25 desalting column (EMP Biotech). Purified protein was concentrated to <500 μl in a centrifugal concentrator with either 30 or 50 kDa molecular weight cut-off (Sartorius VivaSpin).

### Protein characterisation

UV/Vis spectrophotometry was performed on a Cary 60 instrument (Agilent). Size exclusion chromatography was performed in PB buffer plus 0.24% Cymal-5 using a 10/300 GL column at a flow rate of 0.5 ml/min. Circular dichroism used a Jasco J-1500 instrument at protein concentrations of 0.2 – 0.5 mg/ml in a 1 mm pathlength cell, and wavelengths at which the HT voltage exceeded 600 V were excluded. Redox potentials were determined as described^28^ in Membrane Buffer including the mediators phenazine ethosulfate (PES), duroquinone (DQ), indigotrisulfonate (ITS), 2-hydroxy-1,4-napthoquinone (2H14NQ), phenazine (PHE) and anthroquinone-2-sulfonate (AQS) plus 0.08 % Cymal-5 buffer (pH 7.4). This detergent concentration was beneath the critical micelle concentration, but was empirically found to sustain the protein-detergent complex while avoiding the partitioning of mediators into empty micelles. Data were fit to the 2-electron Nernst equation. For confocal microscopy, cells were fixed with 2% paraformaldehyde in PBS and cured on glass slides with ProLong Gold antifade reagent (ThermoFisher). Images were collected with a Leica SP8 AOBS confocal laser scanning microscope and analysed with Fiji^29^.

For native mass spectrometry, solutions containing purified CytbX were buffer-exchanged into 0.2 M ammonium acetate (pH 8.0) containing 2x CMC of Lauryldimethylamine oxide (LDAO) using Biospin 6 columns (Biorad). 2-3 µL of 5 µM protein solution was introduced directly into Q-Exactive UHMR mass spectrometer (ThermoFisher) through gold coated capillary needles that were prepared in-house^30^. The data were collected in positive polarity using the following optimized conditions: Capillary voltage, 1.2 kV; S-lens RF, 200%; Trapping gas pressure, 8.0; Capillary temperature, 200°C; HCD cell, 0 V; In-source trapping, -50 V to -150 V; Resolution, 12,500. For efficient transfer of ionized ions, ion optics were set to injection flatapole, 5 V; inter-flatapole lens, 4 V; bent flatapole, 2 V; transfer multipole, 0 V. The quadrupole was scanned from 1000 to 20,000 *m/z* range. Data were analysed by Xcalibur 4.2 (ThermoFisher) and UniDec^31^ software packages.

To further assess heme loading, CytbX protein concentration was determined with a BCA assay (ThermoFisher 23235) against a standard curve generated from a non-heme *de novo* membrane protein^16^. The heme concentration of CytbX was determined by comparing heme absorption at 560 nm of the as-purified protein, which was assumed to be fully-oxidised, and the same sample after reduction with a few grains of dithionite. The difference in extinction coefficient between the oxidised and reduced forms was taken as 22,000 M^-1^.cm^-1^ as described^32^. Alternatively, CytbX was treated with 0.5 M NaOH and 10% pyridine, and the hemochrome concentration determined using ?_557_ of 28,150 M^-^1.cm^-1^ as described ^33^. These two methods were in close agreement and supported the 2:1 heme:protein ratio determined by mass spectrometry. This also allowed the calculation of extinction coefficients for diheme CytbX in the oxidised state (ε_418_ = 155,200 M^-1^.cm^-1^) and the reduced state (ε_432_ = 238,000 M^-1^.cm^-1^).

### Electron transport assays

For interprotein electron transfer 2 μM CytbX, 5 μM *E. coli* flavodoxin reductase (FLDR) and 100μM NADPH were mixed under anaerobic conditions in Membrane Buffer with 0.08% Cymal-5. Absorption spectra were measured every 1.5 s on an Ocean Optics USB2000+ UV-Vis spectrophotometer. Quinone assays were adapted from Lundgren et al^34^, and *E. coli* cytochrome *bo3* ubiquinol oxidase was purified in *n-*dodecyl--D-maltopyranoside^35,36^. For the superoxide reaction, the assay mix was 50 mM sodium phosphate, pH 8, 0.1 mM hypoxanthine, 0.1 mM DTPA, 5U/ml catalase and 0.1 U/ml xanthine oxidase with the addition of 5 μM WST-1, 1.5 μM CytbX, 0.6 μM ubiquinol oxidase and 20 μM Q2 as required. After mixing, scans were immediately collected between 350-600 nm over 30 min at 25°C under aerobic conditions. Approximate % reduction was calculated from the signal ratio at 432/418nm versus dithionite reduction. For the reverse reaction, reduced quinol (Q2H_2_) was prepared using sodium borohydride^34^ and used immediately with or without 0.1 mM WST-1.

## RESULTS

### Membrane protein design

The exterior surface of soluble 4D2 was computationally redesigned with a limited alphabet of hydrophobic and small polar residues (FAILVWGST) using RosettaMP with the *franklin2019* energy function (Fig. 1a). In preliminary work we found that allowing the fullest degree of design freedom over the remainder of the sequence did give rise to biocompatible membrane proteins, but that these designs did not clearly show diheme binding during recombinant expression. To overcome this, we adopted a more conservative strategy which explicitly preserved the interior packing residues responsible for specific knobs-into-holes interactions at coiled-coil register positions *a* and *d*, as well as threonines occurring at four *g* positions that form ‘keystone’ hydrogen bonds to the heme-coodinating histidines^18^. We also retained 19 other residues from 4D2 that made crucial heme contacts (including the four axial histidines), the N-terminal Gly and Ser, and four proline helix caps. Soluble interhelical loops were also preserved during design, except that Val 57 of 4D2 was specified as either Lys or Arg to dictate transmembrane topology according to the positive inside rule. The positions of both designable and conserved residues are shown in Fig. 1a and 1b.

**Figure 1:**
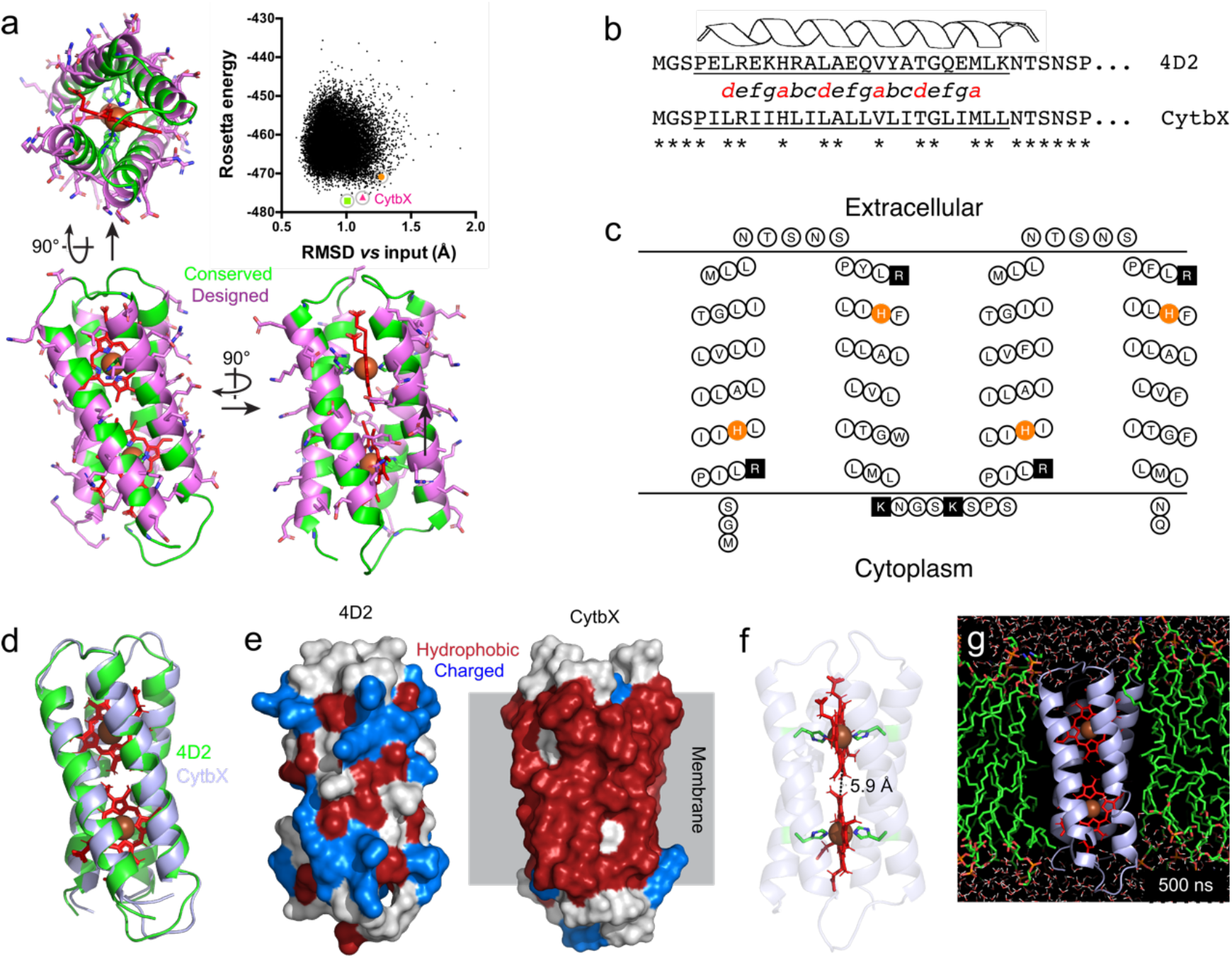
Design of CytbX from the soluble *de novo* protein 4D2. (**a**) The RosettaMP design suite was used to mutate designated surface residues of 4D2 (*magenta*). The subsequent energy *vs* RMSD plot for over 18,000 decoys was tightly clustered. The construct subsequently termed CytbX (*pink triangle*) was one of three low-scoring decoys selected for further study. (**b**) CytbX shows substantial sequence variation from the 4D2 parent at designable positions, with Helix 1 given as an example. The helical register shows the critical coiled-coil packing positions *a* and *d* in red. (**c**) Expected transmembrane topology of CytbX. Histidine residues for heme coordination and positively-charged residues, including the loop lysines, are highlighted *orange* and *black* respectively. Figure drawn with TOPO2 (http://www.sacs.ucsf.edu/TOPO2/). (**d**) The parent 4D2 fold is retained in CytbX, with RMSD of 0.5 Å between the two structures. (**e**) The design process results in a hydrophobic protein surface compatible with a membrane interior. (**f**) The CytbX fold supports the bis-histidine coordination of two heme cofactors with an edge-to-edge distance of approximately 6 Å. (**g**) Snapshot of CytbX after 500 ns molecular dynamics simulation in an *E. coli* lipid bilayer.

All 18,400 of the design decoys were tightly clustered with Rosetta energy scores of -440 to -480, with an average RMSD of about 1 Å from the 4D2 parent structure (Fig. 1a). Three decoys were selected for further analysis and characterisation based upon their overall score and packing statistics (Fig. 1a). In recombinant expression trials one of these variants was produced at substantially higher yields than the other two (see below and Supplementary Figure S3). This construct was the focus of all further analysis and was termed Cytochrome *b*X (CytbX). Bioinformatic analyses and *ab initio* structure prediction suggested that CytbX would indeed form a four-helix transmembrane bundle as designed (Supplementary Figure S4).

The primary structure of CytbX was significantly changed from 4D2, with 49 of 113 amino acid positions altered from the input sequence (Fig. 1b and 1c), whereas the overall structure was essentially unchanged (Fig. 1d). Because of the limited design alphabet and the requirement for membrane compatibility, the surface of CytbX was dominated by hydrophobic residues (Fig. 1e) and in particular Leu and Ile which constituted 44% of all residues. A BLASTP search of the non-redundant NCBI database confirmed that there was no sequence homology between CytbX and any natural protein. The two hemes coordinated by CytbX were retained in the same conformation as in 4D2 (Fig. 1f), with edge-to-edge distances of approximately 6 Å compatible with rapid electron transport between the heme groups^37^. Figure 1g shows the structure of CytbX after 500 ns of a molecular dynamics simulation in a 3:1 DOPE:DOPG lipid bilayer. Some simulations showed a slight straightening of helical packing angles between two adjacent helices from about 150° to about 165° but otherwise both the apo-and holoprotein models of CytbX were stable over the simulation timecourse, with limited fluctuation of the transmembrane helices (Supplementary Figure S5).

### Cellular production of CytbX

A synthetic gene encoding CytbX was expressed from a recombinant plasmid in *Escherichia coli* strain C43(DE3), a strain that was previously selected for membrane protein production^26^. CytbX was well-tolerated by the cells, with no growth inhibition upon induction. Confocal fluorescence microscopy confirmed the successful localisation of a CytbX-GFP fusion protein to the bacterial membrane (Fig. 2a), suggesting that the first transmembrane helix of CytbX can act as an intrinsic cotranslational signal sequence. Cell cultures were supplemented with the heme precursor *8*-aminolevulinic acid (ALA) at the point of induction to upregulate heme biosynthesis and enhance incorporation into CytbX, and the resulting cell pellets were bright red (Fig, 2b) indicating the presence of a new cellular heme sink. When cells were lysed and fractionated, this red colouration was strongly associated with cellular membranes.

**Figure 2:**
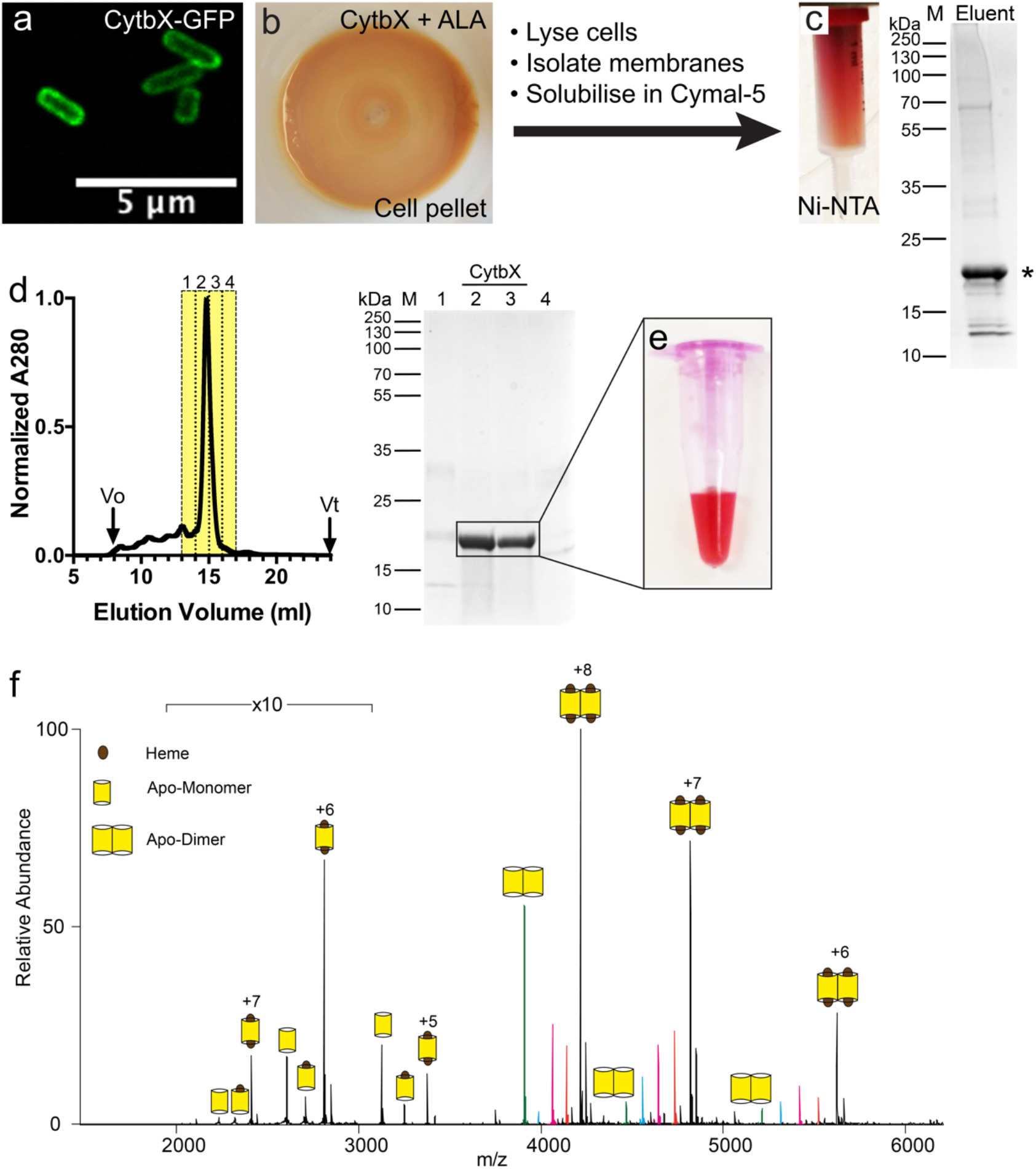
Cellular production and purification of CytbX. (**a**) CytbX-GFP fusion protein is localised to the cytoplasmic membrane of *E. coli*. (**b**) *E. coli* strains overexpressing CytbX turn red when the culture is supplemented with the heme precursor ALA. (**c**) CytbX can be partially purified by immobilized metal affinity chromatography. (**d**) Semi-purified CytbX is purified to homogeneity using size-exclusion chromatography. The major peak on the chromatogram is confirmed as CytbX by SDS-PAGE with Coomassie staining. (**e**) Purified CytbX is coloured red, consistent with heme binding. Image shows protein as-purified from the cell without addition of any exogenous cofactor. (**f**) Native mass spectrometry reveals that CytbX forms a persistent dimer that coordinates two hemes. Coloured peaks within the dimeric charge series correspond to the number of hemes bound per dimer. The major dimer population contains four hemes (*black*) with minor populations corresponding to three (*orange*), two (*magenta*), one (*cyan*) and no heme (*green*).

### Purification of CytbX

Cellular membranes containing His_10_-tagged CytbX were solubilised with the gentle nonionic detergent Cymal-5 prior to immobilised metal affinity chromatography (IMAC). The protein-detergent complex retained a bright red colour, was resolved as a single peak on size-exclusion chromatography (SEC), and could be purified to homogeneity (Fig. 2c-e). No additional heme was added at any stage of the preparation. Yields were typically 0.3 mg of CytbX per litre of original bacterial culture. CytbX migrated slightly slower than the theoretical molecular weight of 15.6 kDa on an SDS-PAGE gel, which is common for integral membrane proteins; the correction factor of 1.13 suggested by Rath and Deber^38^ adjusted the apparent weight to the theoretical weight. Native mass spectrometry (Fig. 2f) revealed that CytbX was purified almost entirely as a dimer, consistent with the SEC elution volume. This CytbX dimer was nearly fully loaded with two hemes per protomer, as determined by the absolute mass of the main peak (33,730 Da versus hypothetical mass of 33,672 Da for the diheme dimer) and by the mass shift of 2460 Da over a small proportion of apoprotein and some minor peaks corresponding to partial heme binding states. This 2:1 heme:protein ratio was corroborated by independent measurements of the protein and heme concentrations in purified samples.

### Characterisation of CytbX

Circular dichroism (CD) spectroscopy of purified CytbX confirmed the expected helicity of this design (Fig 3a). CytbX was highly thermostable, showing only marginal changes in secondary structure and no aggregation at temperatures up to 95°C (Fig. 3a). The visible absorption spectra of CytbX were consistent with the loading of two *b*-type hemes per protein and were indistinguishable from natural membrane cytochromes (Fig. 3b). The as-purified protein contained oxidised heme, with a Soret band at 418 nm. CytbX could be rapidly reduced with dithionite, causing a shift in the Soret band to 430 nm and the emergence of sharp β and α bands at 532 nm and 561 nm respectively. Redox potentiometry (Fig. 3c) revealed clear redox splitting of the hemes with two midpoint potentials (*E*_*m*_) at -14 and -127 mV *vs*. NHE at pH 7.4. We did not observe the redox hysteresis of the original D2 peptide assembly^18^, and the interheme splitting of 113 mV for CytbX is nearly double that of the 4D2 parent^17^. The two heme chromophores of CytbX should be close enough to exhibit exciton coupling, and this was confirmed by visible CD (Fig. 3d).

**Figure 3.**
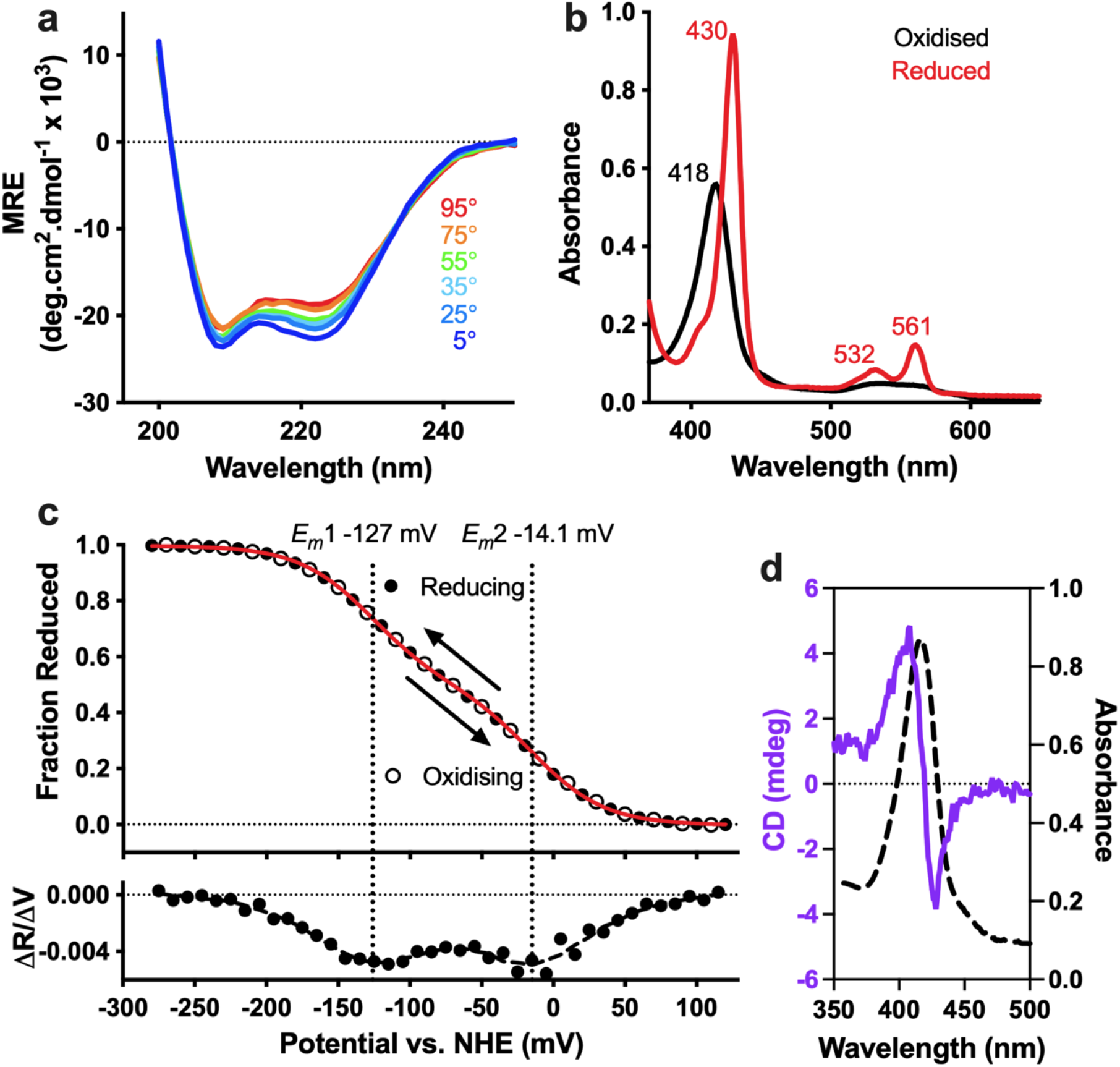
Characterisation of CytbX. (**a**) CytbX is strongly alpha-helical and resists thermal unfolding, with only a marginal change in circular dichroism spectra up to 95°C. *MRE*, mean residue ellipticity. (**b**) UV-Vis absorption spectra of as-purified CytbX in the oxidised and the dithionite-reduced form at room temperature. (**c**) Redox potentiometry is consistent with the binding of two hemes with split potentials. *Red line*, fit to the 2-electron Nernst equation. *Lower panel* shows the first derivative, with smoothed data as a dashed line. (**d**) The proximity of the two hemes leads to exciton coupling in visible circular dichroism.

### CytbX in model interprotein electron transport chains

CytbX was next explored as the terminal acceptor in a minimal electron transport chain. This was based upon a soluble redox protein partner, the *E. coli* flavodoxin-NADP_+_ oxidoreductase (FLDR). FLDR, which has an *E*_*m*_ of -288 mV, normally mediates the transfer of electrons from NADPH (*E*_*m*_ of -372 mV) to flavodoxin^39^ but can directly reduce some other proteins *in vitro*^40^. The NADPH/FLDR pair was able to rapidly and fully reduce purified CytbX under anaerobic conditions (Fig. 4a-d). This confirms that the membrane-embedded heme centres of CytbX are accessible to diffusive redox partners and that elementary electron transport chains can be constructed linking biologically relevant, obligate 2-electron donors to CytbX without the need to design specific protein-protein interactions.

**Figure 4:**
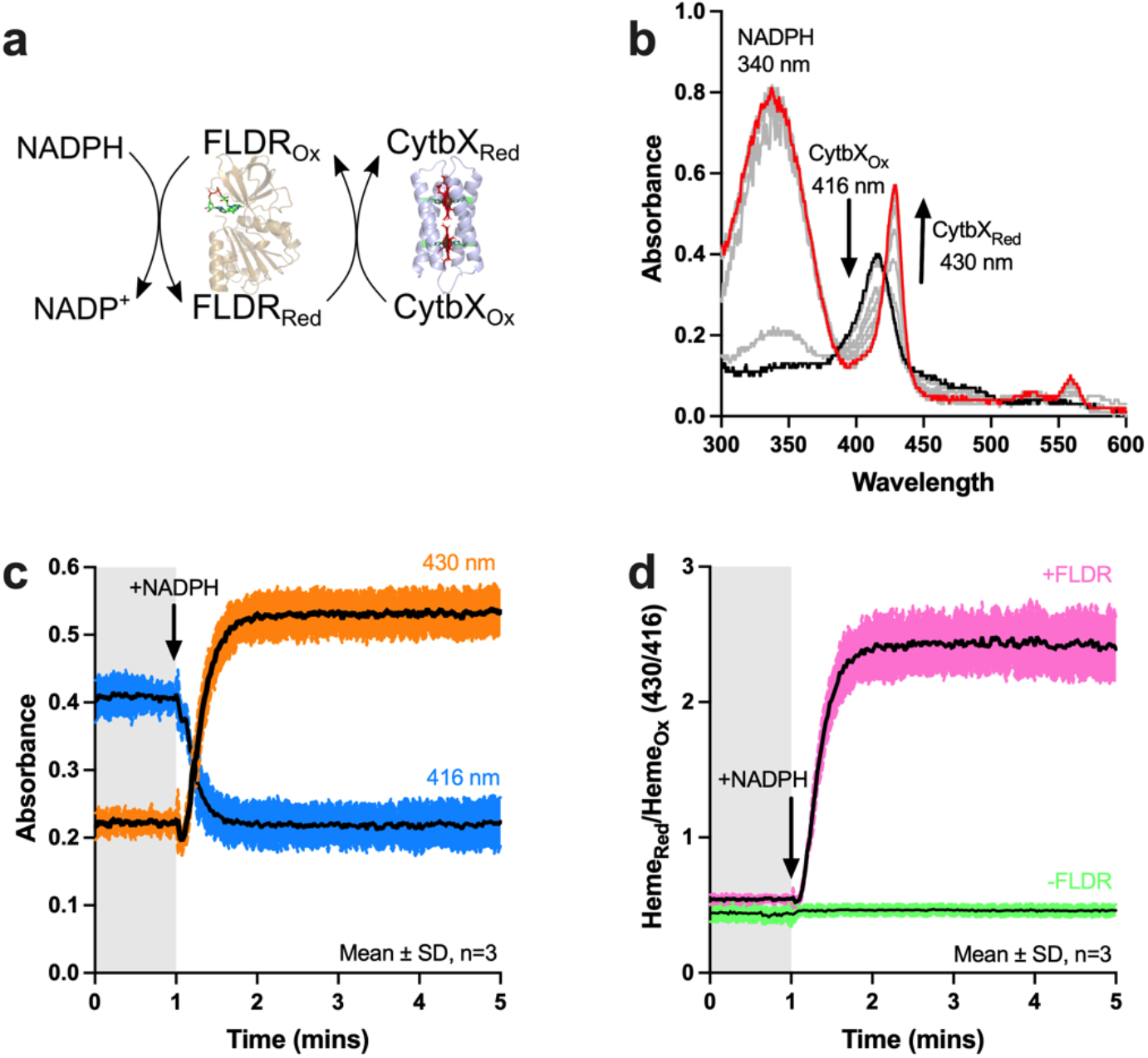
CytbX as the terminal acceptor in an electron transport chain. (**a**) Purified *E. coli* flavodoxin reductase (FLDR) mediates the transfer of electrons from NADPH to CytbX. FLDR structure is from PDB 1FDR. (**b**) The reaction proceeds immediately upon addition of NADPH (*black line*) and CytbX is fully reduced after 1 min (*red line*). Intermediate times are shown in grey, with NADPH maintained in excess throughout. (**c**) The appearance of the reduced Soret band at 430 nm is simultaneous with the loss of the oxidised band at approximately 416 nm. (**d**) Data transformed from panel (**c**) compared with an equivalent control omitting FLDR.

### CytbX in quinone-based electron transport chains

One of the core functions of natural membrane cytochromes is electron transfer reactions with membrane-soluble quinones. To explore whether CytbX had any intrinsic quinone reactivity to quinones, methods were adapted from the study of *E. coli* CybB, a small diheme four-helix bundle which is a membrane superoxide:quinone oxidoreductase^34,41^. An overview of this assay system is depicted schematically in Fig. 5a. The production of superoxide in these assays via either (i) quinol reduction or (ii) xanthine oxidase (XO) activity is detected spectroscopically through the radical cleavage of dye WST-1 to a stable yellow product with absorption at 445 nm. CybB can compete with WST-1 for superoxide depending upon the presence of quinone as the final electron acceptor^34,42^. The short-chain ubiquinone analogue Q2 is used in these assays because of the extremely low solubility of the natural Q10.

**Fig 5:**
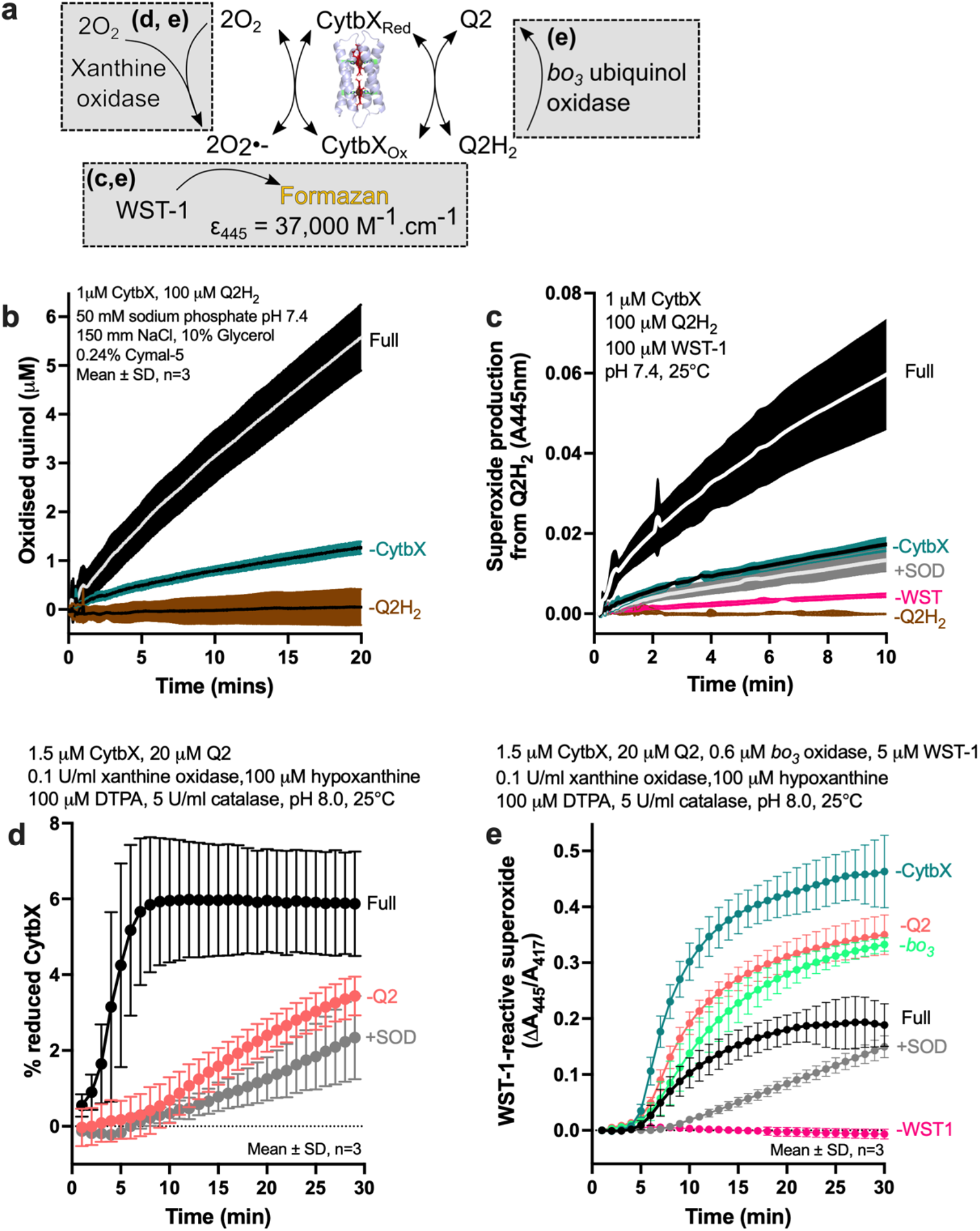
CytbX as a superoxide:ubiquinol oxidoreductase. (**a**) Cartoon schematic of the reconstituted electron transport chain. CytbX couples an enzymatic superoxide-generating system to reduction of the soluble quinone Q2. *E. coli* ubiquinol oxidase (cytochrome *bo*_*3*_) regenerates the quinone pool. Reactions highlighted in grey boxes are relevant to other panels as shown. (**b**) CytbX oxidises ubiquinol Q2H_2,_ determined by Q2 absorption at 275 nm. (**c**) Quinol oxidation by CytbX simultaneously produces superoxide, detected by radical cleavage of WST-1. Controls include the inclusion of superoxide dismutase (*+SOD*). (**d**) Enzymatically-generated superoxide reduces CytbX, as determined by changes in heme spectra. A persistent reduced state is formed in the presence of quinone/quinol, see text for discussion. (**e**) Reconstitution of the full reaction shown in Panel (**a**). CytbX competes with WST-1 for superoxide produced by XO, with competition being most effective when the soluble quinone pool is replenished via *bo3* ubiquinol oxidase (*filled black circles*).

A first set of experiments assessed whether CytbX could catalyse the oxidation of quinol Q2H_2_. These experiments monitored the production of the oxidised product, quinone, by measuring the quinone absorption band at 275 nm. Quinol oxidation by CytbX was clearly observed above background levels over a timecourse of several minutes (Fig. 5b). The subsequent instantaneous oxidation of quinol-reduced CytbX resulted in detectable levels of superoxide (Fig. 5c). CytbX thus exhibits quinol oxidase activity, although the rate of this reaction is below that of the natural enzyme CybB^34^.

CytbX was next employed in the reverse reaction to catalyse the relay of electrons from enzymatically-generated superoxide to quinone Q2. Unlike CybB, CytbX does not form a persistent reduced state in the presence of superoxide (Fig. 5d). However, when Q2 was also included in the reaction, a small proportion of CytbX was found to be reduced as superoxide was generated. This suggests that CytbX uses electrons from superoxide to mediate the formation of Q2H_2_ from Q2, with the combined reducing power of both superoxide and Q2H_2_ then leading to a persistent fraction of reduced heme.

Finally, the full assay was reconstituted in which CytbX competes for superoxide with WST-1 (Fig. 5e). This competition was most pronounced when cytochrome *bo3* ubiquinol oxidase was used to maintain an excess of oxidised quinone, as seen for CybB^34,42^. The extent of competition was much lower for CytbX than that observed for CybB, consistent with the evolutionary specialisation of that enzyme versus the intrinsic activity of CytbX.

## Discussion

The computational design of integral membrane proteins has achieved specific transmembrane folds^43-46^ and functions such as metal transport^47^ and ion permeation^48^. The current study now expands this effort by introducing a recombinant construct, CytbX, that is capable of cofactor recruitment in cellular membranes. The results, which are strongly consistent with the designed model, show that this *de novo* membrane hemoprotein can recapitulate features associated with natural respiratory and photosynthetic complexes such as the splitting of heme redox potentials. The metal centres of CytbX can engage in chemical, electrochemical and biochemical electron transfer even though such functions were not explicitly considered during design.

The successful cellular production of CytbX confirms that, in principle, *de novo* membrane proteins are compatible with cellular lipid bilayers. Nonetheless -just as for natural membrane proteins -not all recombinant sequences are tolerated equally by the cell, and expression screening is needed to identify variants that express at higher levels. Our working assumption is that CytbX is trafficked and inserted into the plasma membrane cotranslationally via classical translocation pathways, with the first transmembrane helix acting as the localisation and integration signal^49^. This remains to be confirmed but would agree with a model in which an apoprotein intermediate of CytbX assembles in the membrane first, and the cofactor is then captured by this pre-existing structure to complete protein folding^50,51^. We have not yet been able to test this idea because of the difficulty in purifying apo-CytbX; while complete heme loading seems to require the addition of the heme precursor ALA, partially-loaded holoprotein is still produced in unsupplemented cultures. Our data are consistent with the concept that cellular heme is maintained in an exchangeable reservoir via weak non-specific protein-heme interactions, and then can pass from this heme pool to tight-binding metalloproteins (which must include integral membrane proteins)^52^. CytbX can apparently compete for the cellular heme pool under normal conditions, with easier access to this cofactor pool when heme levels are elevated by the addition of ALA. Further work might involve the iterative redesign of CytbX for tighter heme binding, in order to achieve full cofactor loading in unsupplemented cells. Overall, the ability of CytbX to recruit cellular heme augurs well for the further design of membrane-embedded metal centres.

The midpoint potentials of CytbX, and the separation between them, are broadly similar to several natural *b*-type diheme membrane cytochromes^2,53-56^. However, heme *b*_*H*_ of the *bc1* complex typically has a positive redox potential^2^ and the values obtained here for CytbX are more negative than some other membrane diheme centres^57^ including the cyt*b*_561_ family of small cytochromes^58-60^. There thus appears to be scope for redox engineering of CytbX to access different potentials. The protein milieu around both heme cofactors is rather similar (though not identical) and so the splitting of heme redox potentials in CytbX is likely to arise from direct interactions between the two cofactors, as proposed by Wikstrom^61^ and observed previously in other *de novo* heme-containing proteins^62^. The separation of these potentials is more pronounced than observed for the soluble 4D2 parent^17^ probably because of the low dielectric of the micelle interior. The redox splitting could also be influenced by the presence of two lysines in intracellular loop 2 (Fig. 1c) which may contribute to a different electrostatic environment at one of the heme sites.

CytbX is an oxidoreductase that can engage in electron transfer reactions (Figs. 4 and 5) even though these activities were not targeted as part of the design. Enhancing the intrinsic functionality of CytbX could be accomplished through coupling to other redox modules via specific protein-protein interaction motifs or gene fusion. This could also be achieved by the binding of alternative and unnatural metalloporphyrins^63^, which can markedly adjust the redox potentials of *de novo* proteins and be a route to functions such as light-induced charge separation^64,65^. This approach can be extended to the incorporation of other redox centres including flavins^65^, iron-sulfur clusters^66,67^, dimetal sites^19,68^ and chemical chromophores^65^. Additionally, it should be possible to design specific small-molecule binding sites, for example to extend the inherent reactivity of CytbX with ubiquinol. Because CytbX is genetically-encoded, this computational redesign could be combined with directed evolution methods that are able to select for protein functionality within the cell.

We thus report the computational surface-swapping of a parametrised helical bundle to produce a *de novo* diheme membrane cytochrome. The functional integration of such artificial membrane cytochromes with living systems should enable the future assembly of novel electron transport complexes and redox catalysts.

## Supporting information

Supplementary data - Rosetta design files

## Acknowledgements

We thank Prof. Christoph von Ballmoos for the kind gift of plasmid pETcyoII, used for the heterologous expression of *E. coli bo3* ubiquinol oxidase. This work was supported by a SynBioCDT doctoral training award to BJH, BrisSynBio, a BBSRC/EPSRC Synthetic Biology Research Centre (BB/L01386X/1) and BBSRC grant BB/W003449/1 to JLRA. This work used the computational facilities of the Advanced Computing Research Centre at the University of Bristol (http://www.bristol.ac.uk/acrc).

## Notes

### Competing Interest Statement

The authors have declared no competing interest.

