## Supplementary Information for "Cellular production of a *de novo* membrane cytochrome"

**Contents**

**Supplementary methods**

**Figure S1**: Comparison of surface-swapped amino acid sequences produced by Rosetta design.

**Figure S2**: Synthetic genes corresponding to each protein design.

**Figure S3**. Purification of three Rosetta designs.

**Figure S4**. Topology prediction and *ab initio* structure prediction of CytbX.

**Figure S5.** Molecular dynamics simulations of CytbX in an explicit bilayer.

**Supplementary Methods**

***Molecular Dynamics simulations***

Relaxed CytbX model structures were inserted into 3:1 DOPE:DOPG lipid bilayers using PACKMOL-Memgen^1^ with bis-histidine coordination of *b*-type hemes as described^2^. The systems were protonated using Reduce, and solvated and parameterised using LEaP. The ff14SB forcefield was used for protein, Lipid17 for lipids, and tip3p for water and ions^3,4^. All molecular dynamics simulations were run using the Amber18 software package^5,6^ with GPU acceleration^7^. Systems were energy minimized by three rounds of 5000 steps steepest-descent minimization, with harmonic restraints for protein non-hydrogen atoms only, for protein C-alpha atoms only and with no restraints, respectively. Systems were then heated to 303K over 100 ps using Langevin dynamics and 1 Bar pressure was then applied using a Berendsen barostat^8^ with semi-isotropic pressure coupling over a further 100 ps (2 ps pressure relaxation). Using GPU acceleration, systems were further equilibrated for 100 ps at 1 Bar and 303 K with restraints to only the C-alpha atoms of the protein. Triplicate unrestrained MD simulations were run for 500 ns each. All water molecules were kept rigid using SETTLE^9^. Trajectories were analysed using CPPTRAJ^10^ and visualised using VMD^11^.

1	Schott-Verdugo, S. & Gohlke, H. PACKMOL-Memgen: A Simple-To-Use, Generalized Workflow for Membrane-Protein-Lipid-Bilayer System Building. *J Chem Inf Model* **59**, 2522-2528, (2019).

2	Yang, L. *et al.* Data for molecular dynamics simulations of B-type cytochrome c oxidase with the Amber force field. *Data Brief* **8**, 1209-1214, (2016).

3	Maier, J. A. *et al.* ff14SB: Improving the Accuracy of Protein Side Chain and Backbone Parameters from ff99SB. *J Chem Theory Comput* **11**, 3696-3713, (2015).

4	MacKerell, A. D. *et al.* All-atom empirical potential for molecular modeling and dynamics studies of proteins. *J Phys Chem B* **102**, 3586-3616, (1998).

5	Case, D. A. *et al.* The Amber biomolecular simulation programs. *J Comput Chem* **26**, 1668-1688, (2005).

6	Salomon-Ferrer, R., Case, D. A. & Walker, R. C. An overview of the Amber biomolecular simulation package. *Wires Comput Mol Sci* **3**, 198-210, (2013).

7	Salomon-Ferrer, R., Gotz, A. W., Poole, D., Le Grand, S. & Walker, R. C. Routine Microsecond Molecular Dynamics Simulations with AMBER on GPUs. 2. Explicit Solvent Particle Mesh Ewald. *J Chem Theory Comput* **9**, 3878-3888, (2013).

8	Berendsen, H. J. C., Postma, J. P. M., Vangunsteren, W. F., Dinola, A. & Haak, J. R. Molecular-Dynamics with Coupling to an External Bath. *J Chem Phys* **81**, 3684-3690, (1984).

9	Miyamoto, S. & Kollman, P. A. Settle - an Analytical Version of the Shake and Rattle Algorithm for Rigid Water Models. *Journal of Computational Chemistry* **13**, 952-962, (1992).

10	Roe, D. R. & Cheatham, T. E. PTRAJ and CPPTRAJ: Software for Processing and Analysis of Molecular Dynamics Trajectory Data. *Journal of Chemical Theory and Computation* **9**, 3084-3095, (2013).

11	Humphrey, W., Dalke, A. & Schulten, K. VMD: Visual molecular dynamics. *J Mol Graph Model* **14**, 33-38, (1996).

**Figure S1**: **Comparison of surface-swapped amino acid sequences produced by Rosetta design.** Sequences are identified by their arbitrary Rosetta simulation ID and aligned in ClustalOmega. 50-289 is renamed CytbX in the accompanying paper.

```
50_289      MGSPILRIIHLILALLVLITGLIMLLNTSNSPYLRLIHFLLALLVLITGWLMLKNGSKSP	60
49_13       MGSPFLRLIHLILALLVLLTGLIMLLNTSNSPYLRLIHFLLAILVLITGLIMLKNGSRSP	60
30_82       MGSPWLRLLHLFLALLVLLTGLIMLLNTSNSPYLRLIHFLLAILVLITGLLMLKNGSRSP	60
            **** **::**:******:***********************:****** :******:**

50_289      SPILRLIHIILAILVFITGIIMLLNTSNSPFLRILHFILALLVFITGFLMLNQ	113
49_13       SPILRLLHLILAILVFLTGLIMLWNTSNSPYLRLIHFLLALLVFLTGFLMLNQ	113
30_82       SPLLRLLHLILAILVFLTGLIMLLNTSNSPYLRLLHLILALLVFITGFLMLNQ	113
            **:***:*:*******:**:*** ******:**::*::******:********
```

**Figure S2: Synthetic genes corresponding to each protein design.** The translation of each gene is shown directly underneath.

>CytbX_synthetic_gene

**ATG**GGCTCTCCTATTCTGCGCATCATTCACCTGATTTTGGCCTTGCTGGTTCTGATTACCGGACTTATCATGCTGCTGAATACGTCAAATAGCCCCTATCTTCGCCTCATTCATTTTTTACTGGCACTGCTCGTGCTGATTACCGGTTGGCTGATGCTAAAAAACGGTAGTAAGAGTCCGAGCCCGATCCTCCGTTTAATCCACATAATTCTGGCAATACTGGTATTTATTACTGGCATCATTATGTTACTGAACACATCGAACAGCCCATTCCTGCGGATTTTGCATTTCATCCTTGCGTTATTGGTCTTTATCACGGGCTTCCTTATGCTGAACCAGGCGGCCGCAGGTAAACCGATCCCGAATCCACTGTTAGGGCTGGATTCCACCCATCACCACCATCACCATCACCATCATCAT**TGA**

>CytbX_translated

MGSPILRIIHLILALLVLITGLIMLLNTSNSPYLRLIHFLLALLVLITGWLMLKNGSKSPSPILRLIHIILAILVFITGIIMLLNTSNSPFLRILHFILALLVFITGFLMLNQAAAGKPIPNPLLGLDSTHHHHHHHHHH*

>CytbX-GFP_gene

**ATG**GGCTCTCCTATTCTGCGCATCATTCACCTGATTTTGGCCTTGCTGGTTCTGATTACCGGACTTATCATGCTGCTGAATACGTCAAATAGCCCCTATCTTCGCCTCATTCATTTTTTACTGGCACTGCTCGTGCTGATTACCGGTTGGCTGATGCTAAAAAACGGTAGTAAGAGTCCGAGCCCGATCCTCCGTTTAATCCACATAATTCTGGCAATACTGGTATTTATTACTGGCATCATTATGTTACTGAACACATCGAACAGCCCATTCCTGCGGATTTTGCATTTCATCCTTGCGTTATTGGTCTTTATCACGGGCTTCCTTATGCTGAACCAGGCGGCCGCAGGTAAACCGATCCCGAATCCACTGTTAGGGCTGGATTCCACCCTCGAGCTGGTGCCGCGCGGCAGCAGTAAAGGAGAAGAACTTTTCACTGGAGTTGTCCCAATTCTTGTTGAATTAGATGGTGATGTTAATGGGCACAAATTTTCTGTCCGTGGAGAGGGTGAAGGTGATGCTACAAACGGAAAACTCACCCTTAAATTTATTTGCACTACTGGAAAACTACCTGTTCCGTGGCCAACACTTGTCACTACTCTGACCTATGGTGTTCAATGCTTTTCCCGTTATCCGGATCACATGAAACGGCATGACTTTTTCAAGAGTGCCATGCCCGAAGGTTATGTACAGGAACGCACTATATCTTTCAAAGATGACGGGACCTACAAGACGCGTGCTGAAGTCAAGTTTGAAGGTGATACCCTTGTTAATCGTATCGAGTTAAAGGGTATTGATTTTAAAGAAGATGGAAACATTCTTGGACACAAACTGGAGTACAACTTTAACTCACACAATGTATACATCACGGCAGACAAACAAAAGAATGGAATCAAAGCTAACTTCAAAATTCGCCACAACGTTGAAGATGGTTCCGTTCAACTAGCAGACCATTATCAACAAAATACTCCAATTGGCGATGGCCCTGTCCTTTTACCAGACAACCATTACCTGTCGACACAATCTGTCCTTTCGAAAGATCCCAACGAAAAGCGTGACCACATGGTCCTTCTTGAGTTTGTAACTGCTGCTGGGATTACACATGGCATGGATGAGCTCTACAAACTCGAACACCACCACCACCACCACCACCACCACCAC**TGA**

>CytbX-GFP_gene_translated

MGSPILRIIHLILALLVLITGLIMLLNTSNSPYLRLIHFLLALLVLITGWLMLKNGSKSPSPILRLIHIILAILVFITGIIMLLNTSNSPFLRILHFILALLVFITGFLMLNQAAAGKPIPNPLLGLDSTLELVPRGSSKGEELFTGVVPILVELDGDVNGHKFSVRGEGEGDATNGKLTLKFICTTGKLPVPWPTLVTTLTYGVQCFSRYPDHMKRHDFFKSAMPEGYVQERTISFKDDGTYKTRAEVKFEGDTLVNRIELKGIDFKEDGNILGHKLEYNFNSHNVYITADKQKNGIKANFKIRHNVEDGSVQLADHYQQNTPIGDGPVLLPDNHYLSTQSVLSKDPNEKRDHMVLLEFVTAAGITHGMDELYKLEHHHHHHHHHH-

>49_13_synthetic_gene

**ATG**GGGTCGCCTTTCCTGCGCCTGATTCATCTGATCCTCGCTCTGTTGGTGCTGCTTACCGGCCTGATTATGTTACTTAATACGAGCAACAGTCCATATCTCCGGTTGATCCACTTCCTACTGGCCATTCTAGTCCTCATTACCGGCCTGATAATGCTGAAAAATGGTTCACGTAGTCCGAGCCCAATTCTGCGTCTTTTGCACCTGATCTTAGCAATTCTGGTATTTCTGACGGGTTTGATCATGTTATGGAACACTTCCAACTCTCCGTACTTGCGCCTGATTCATTTTCTTTTAGCGCTGTTAGTTTTTCTGACAGGCTTCCTGATGCTGAACCAGGCGGCCGCAGGTAAACCGATCCCGAATCCCCTGCTCGGACTGGATAGCACCCATCATCATCACCATCACCATCATCACCAC**TGA**

>49_13_translated

MGSPFLRLIHLILALLVLLTGLIMLLNTSNSPYLRLIHFLLAILVLITGLIMLKNGSRSPSPILRLLHLILAILVFLTGLIMLWNTSNSPYLRLIHFLLALLVFLTGFLMLNQAAAGKPIPNPLLGLDSTHHHHHHHHHH*

>30_82_synthetic_gene

**ATG**GGCTCTCCGTGGTTACGGCTTCTGCACCTGTTTTTAGCATTATTGGTACTGCTCACCGGCCTCATCATGCTTCTGAACACTAGTAATTCGCCCTACCTGCGCCTGATACACTTCTTGTTAGCTATTCTGGTTTTAATTACGGGTCTGCTGATGCTGAAAAACGGCAGCCGTTCACCGAGCCCGCTCCTACGTCTACTGCACTTGATTCTGGCGATCCTGGTGTTTTTAACGGGGCTTATTATGCTGCTGAACACATCCAATAGTCCATATCTTCGCCTTCTGCATCTGATCCTGGCCCTGCTGGTCTTTATCACCGGTTTCCTGATGCTCAATCAGGCGGCCGCAGGTAAACCGATTCCTAACCCATTGTTGGGATTGGATAGCACCCACCATCACCACCATCATCATCATCATCAT**TGA**

>30_82_translated

MGSPWLRLLHLFLALLVLLTGLIMLLNTSNSPYLRLIHFLLAILVLITGLLMLKNGSRSPSPLLRLLHLILAILVFLTGLIMLLNTSNSPYLRLLHLILALLVFITGFLMLNQAAAGKPIPNPLLGLDSTHHHHHHHHHH*

**Supplementary Figure 3.**

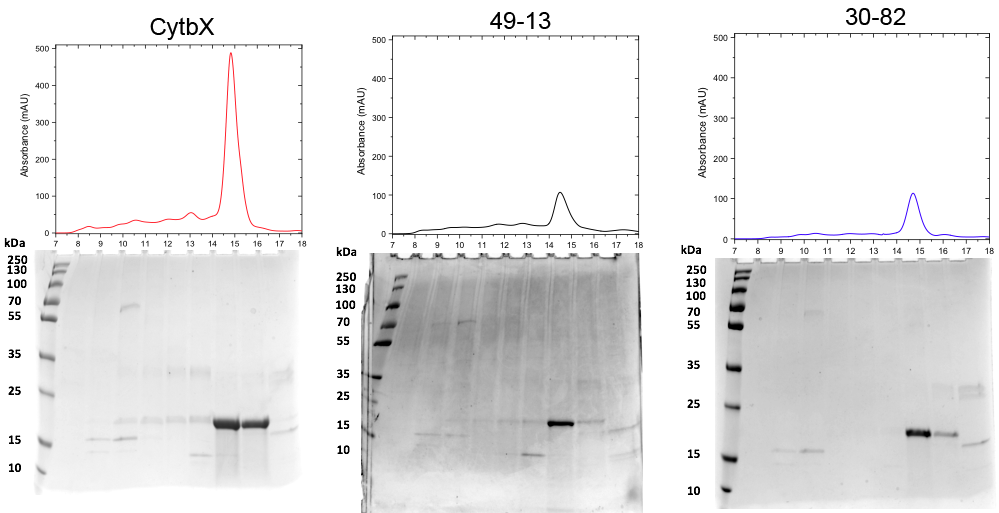

**Figure S3. Purification of three Rosetta designs**. *Top*, size-exclusion chromatography of designs solubilised from *E. coli* membranes in Cymal-5; *Bottom*, Coomassie-stained SDS-PAGE of column fractions. Data for CytbX are reproduced from main text. Two other designs, known only by their Rosetta IDs 49-13 and 30-82, are the green square and orange circle, respectively, on Fig. 1a. Both variants successfully overexpressed and are monodisperse in Cymal-5 but give purification yields ~20% that of CytbX.

**Supplementary Figure 4.**

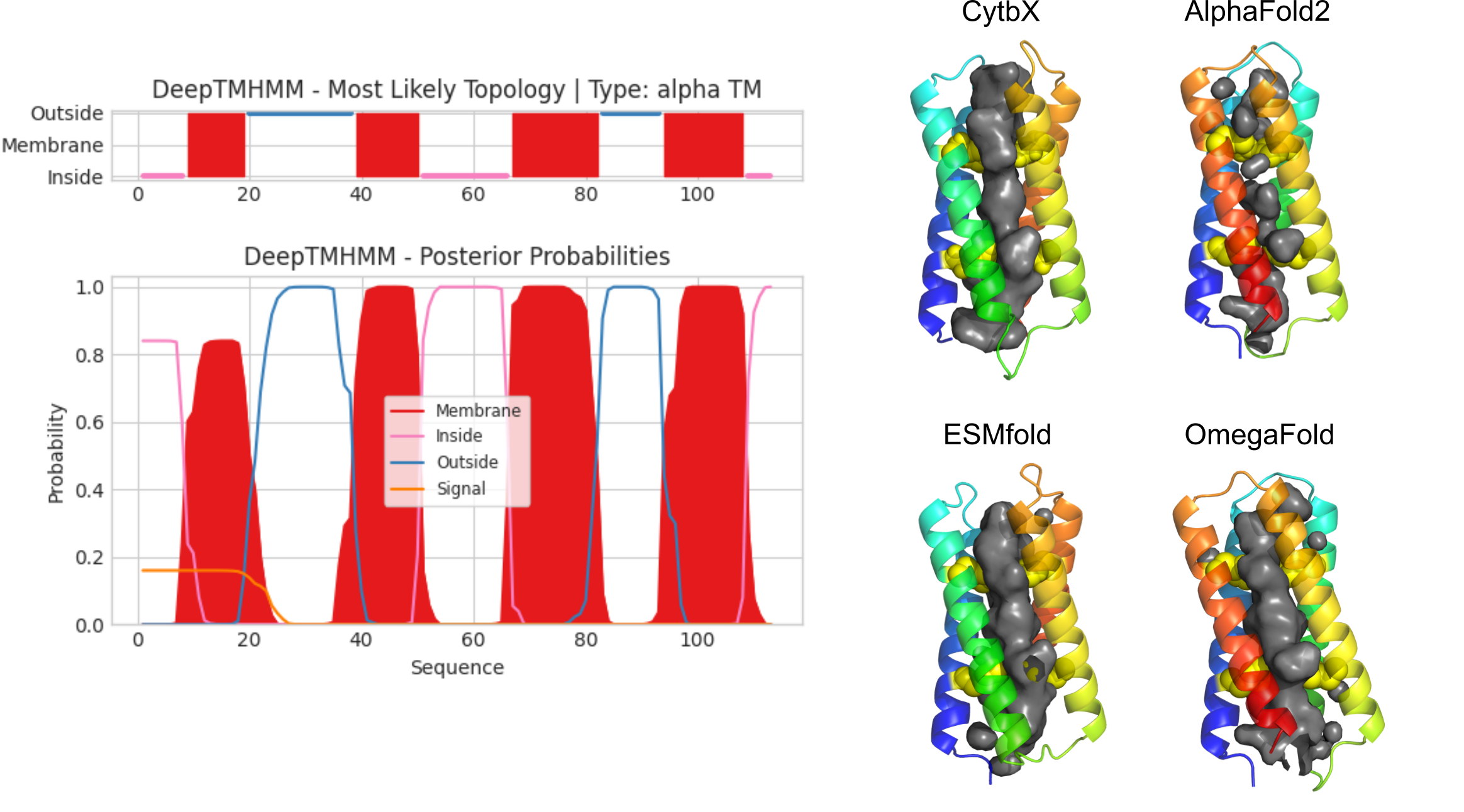

**Figure S4**. Topology prediction and *ab initio* structure prediction of CytbX. *Left panel* shows the results of a DeepTMHMM prediction of transmembrane topology, strongly suggesting that the CytbX sequence will form four transmembrane alpha helices in an N_in_/C_in_ orientation. *Right panels* show CytbX structures as predicted by three different machine learning methods shown. AlphaFold2^1^ predicts an alternative conformation but does not predict an internal cavity for heme binding; this same outcome was also observed for the original 4D2 construct and is at odds with the known experimental structure (PDB 7AH0). In contrast the language-based models ESMfold^2^ and OmegaFold^3^ do predict a heme cavity. ESMfold model differs from the CytbX design by only 0.6 Å.

1	Jumper, J. *et al.* Highly accurate protein structure prediction with AlphaFold. *Nature* **596**, 583-589, (2021).

2	Lin, Z. *et al.* Language models of protein sequences at the scale of evolution enable accurate structure prediction. *bioRxiv*, (2022).

3	Wu, R. *et al.* High resolution *de novo* structure prediction from primary sequence. *bioRxiv*, (2022).

**Supplementary Figure 5.**

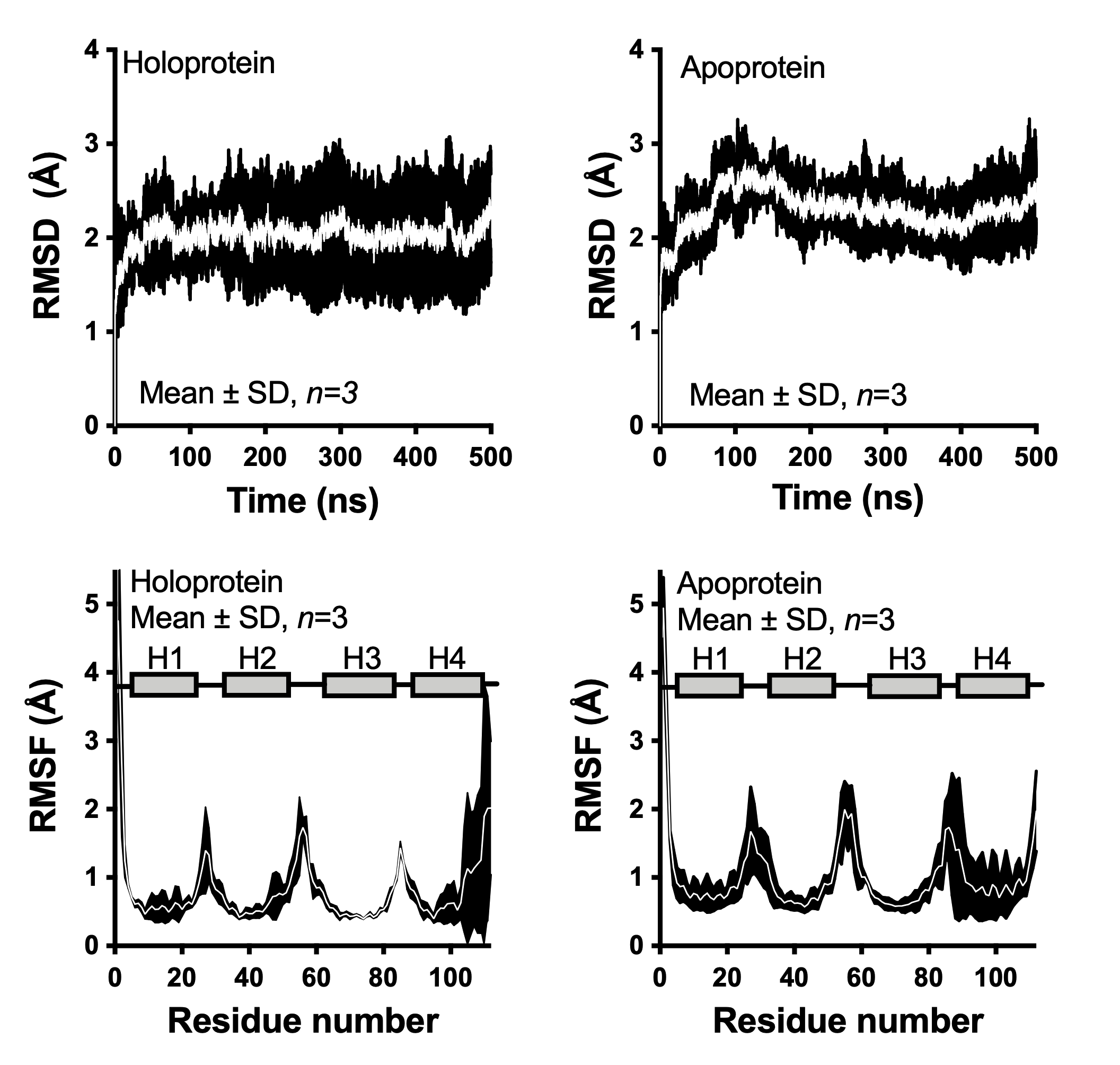

**Figure S5. Molecular dynamics simulations of CytbX in an explicit bilayer.** Cα RMSD plots show that both the apo- and holoprotein equilibrate rapidly and are stable thereafter. Backbone fluctuations are low in the helical regions of the protein, schematised as grey rectangles *H1-4* in the bottom panels. The apoprotein exhibits slightly greater fluctuation overall.
